# Methylation genome wide profiling in lowly and highly efficient adipose somatic cell nuclear transfer in pigs

**DOI:** 10.1101/2022.07.25.501434

**Authors:** Maciej Grzybek, Krzysztof Flisikowski, Tom Giles, Marta Walczak, Rafal Ploski, Piotr Gasperowicz, Richard D. Emes, Pawel Lisowski

## Abstract

Swine is a common model organism for biomedical research. Epigenetic reprogramming in SCNT embryos does not fully recapitulate the natural DNA demethylation events at fertilisation. This study aimed to conduct a genome-wide methylation profiling to detect differentially methylated regions (DMRs) responsible for epigenetic differences in stem cells that displayed high and low efficiency of SCNT and to elucidate the low efficiency of cloning rate in pigs. Adipose tissue mesenchymal stem cells (AMSC)s lines were isolated from adipose tissue of adult male pigs (n=20; high-efficiency cells=10; low efficiency cells= 10). Reduced representation bisulfite sequencing (RRBS) was performed on an Illumina HiSeq1500. Paired-end reads were filtered to remove the adapter contamination, and low-quality reads using TrimGalore!. Filtered reads were mapped to the reference genome using Bismark. MethylKit was used to identify differentially methylated regions (DMRs) (bases and tiles), showing statistically significant differential methylation between two groups: high and low-efficiency AMSCs. Hierarchical cluster analysis according to methylation patterns clearly defined groups with low and high cloning efficiency. We report 3704 bases with statistically significant differences in methylation and 10062 tiles with statistically significant differences in methylation. Most differentially methylated sites are intergenic 62%, 31% are intronic, 4% are located in exons and 4% in promoters. 37% of differentially methylated sites are located in known CpG islands (CGIs) and 4% in CpG island shores (CGSs).

## 1 Introduction

Animal-based disease modeling has become an interest in biomedical research, including cancer, metabolic, cardiovascular and neurological disorders (Groenen et al., 2012a; Arends et al., 2016; Grzybek et al., 2017a; Walters et al., 2017; Schachtschneider et al., 2021).

Swine has been an interest for basic and applied biomedical research for more than 20 years (Rideout et al., 2001; Wilmut et al., 2002; Yang et al., 2007). Swine play essential roles as models of human diseases (Figure 1.), including cardiovascular disease, cancer, diabetes, toxicology and lipoprotein metabolism as a model organism (Bendixen et al., 2010; Zhao et al., 2010; Flisikowska et al., 2013; Walters and Prather, 2013). The first generation of a pig using the SCNT method was conducted in 2000 (Betthauser et al., 2000; Onishi et al., 2000; Polejaeva et al., 2000), and since this time, many genetically modified cloned pigs have been generated (Lai et al., 2002; Lai and Prather, 2003; Li et al., 2006). Despite the success in generating of cloned individuals, there are still limitations that need to be improved to increase the efficiency of the porcine SCNT technique.

**Figure 1.**
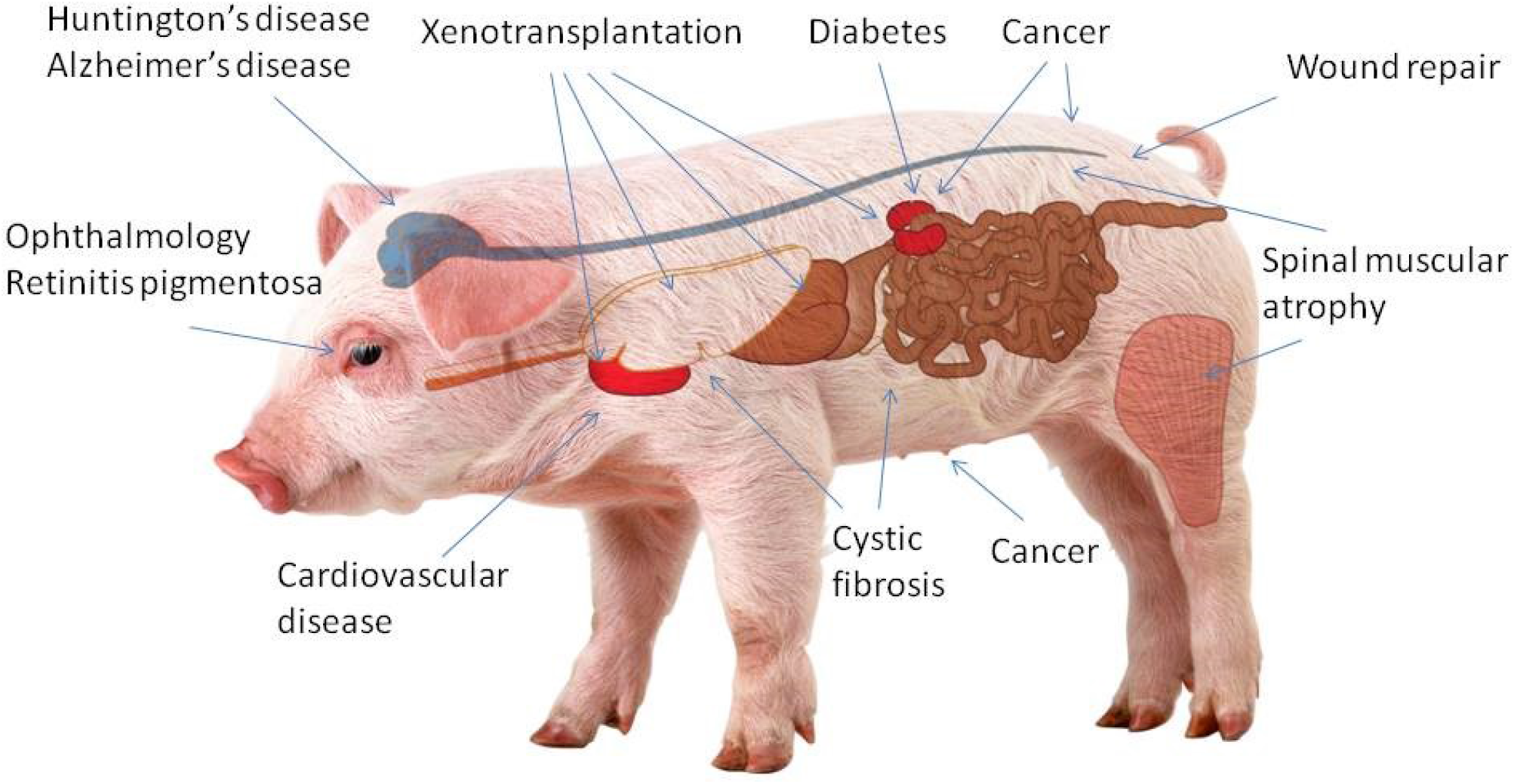
Swine became a model organism for biomedical research due to their similarity to humans. Pigs are ideal organisms to study human health and disease. Their genome is three times closer than the mouse genome to that of humans.

**Figure 2.**
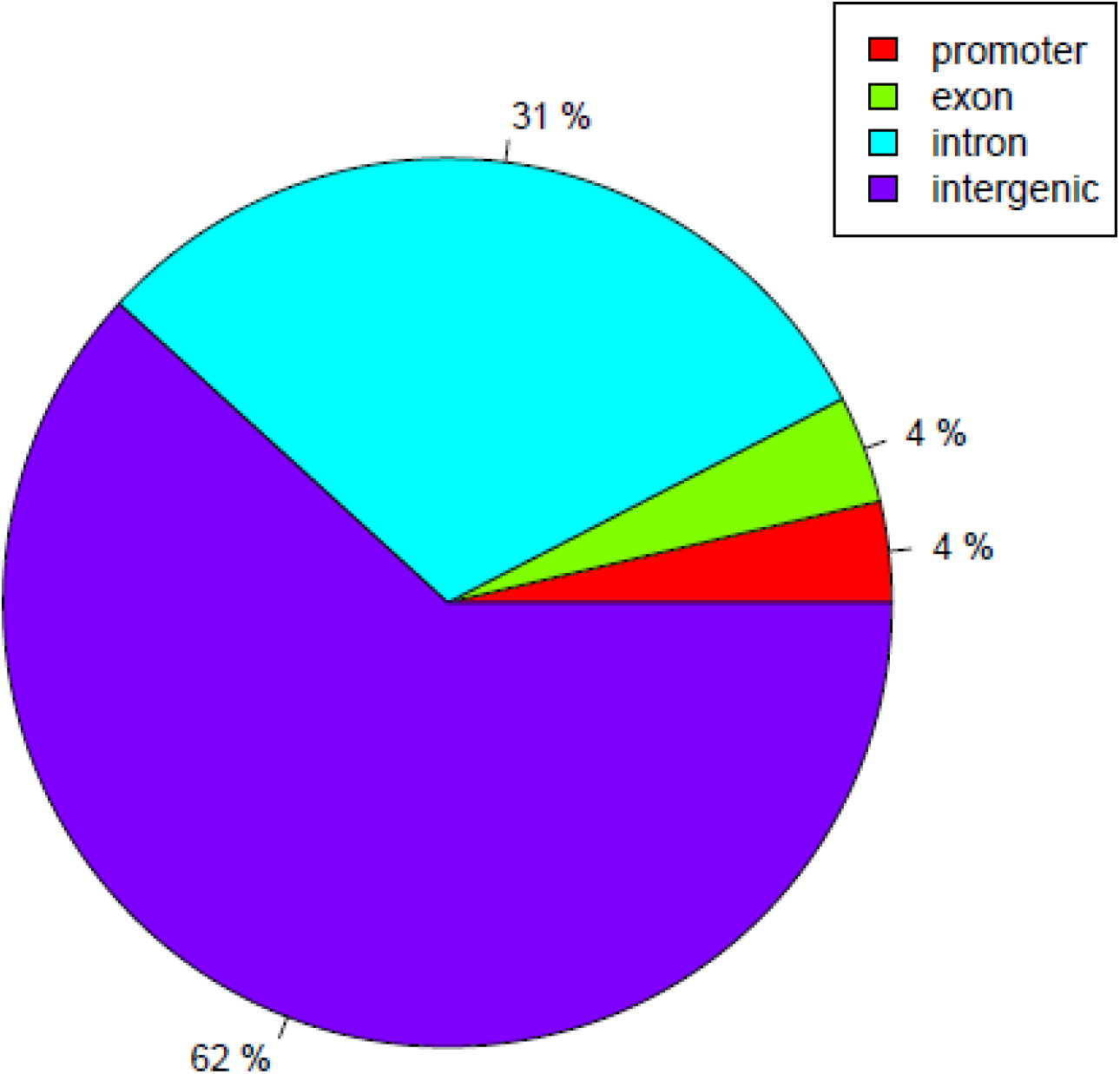
Differential methylation annotations of CpG.

The efficiency of the SCNT method in swine models varies from 0.2% to 7% of newborns per constructed embryo (Fulka and Fulka, 2007; Yang et al., 2007; Zhao et al., 2010). The low success rate limits the extensive application of the pig SCNT technique in biomedical research or agricultural purposes (Kurome et al., 2013). The SCNT method uses somatic cells with low viability and not fully reprogrammed epigenetic memory. This causes swine models to develop malformations (i.e. underweight, cardiac dysfunctions, immunological dysfunctions) (Amiridze et al., 2008; Swindle, 2009; Swindle et al., 2012). Due to the high prevalence of these abnormalities, epigenetic disorders are believed to cause mentioned symptoms during embryo development rather than genetics.

DNA methylation is an essential element in the epigenetic regulation of embryonic development, and it occurs at most CpG dinucleotides in the mammalian genome (Bird, 2002; Fulka and Fulka, 2007; Suzuki and Bird, 2008). Long-term selection and adaption towards high prolificacy and meat production have transformed porcine epigenetics (Li et al., 2012), along with associated genotypic and phenotypic changes (Groenen et al., 2012a; Li et al., 2013) resulting from the modification of the epigenetic regulation of chromatin structure and transcriptional activity. During the transformation process, the porcine DNA methylome displays variable patterns in different breeds and sexes of pigs and variations in other anatomic tissues (Li et al., 2012; Wang and Kadarmideen, 2019). Here we analysed pig methylome of adipose tissue mesenchymal stem cell lines displaying high and low efficiency for live-born piglets in somatic cell nuclear transfer using the RRBS method.

## 2. Material and methods

### 2.1. Ethical approval

All animal experiments were approved by the Government of Upper Bavaria (permit number 55.2-1-54-2532-6-13) and performed according to the German Animal Welfare Act and European Union Normative for Care and Use of Experimental Animals.

### 2.2. Cell lines

Adipose mesenchymal stem cells (AMSC)s lines were isolated from adipose tissue of adult male pigs (n=10 per experimental group) according to standard isolation protocol. Cells were maintained in DMEM supplemented with 10% fetal bovine serum (FBS), 100U/ml of penicillin and 100mg/ml of streptomycin (Invitrogen) at 37°C and 5% CO_2_. The HCT116 DNMT1(2/2) DNMT3b (2/2) double knockout clone number 2 (DKO) cell line was a kind gift from Dr Steve Baylin. The cell line was grown in McCoys’5A medium with 10% FBS, 0.2mg/ml Neomycin, and 0.1mg/ml. Genomic DNA was extracted using standard phenol: chloroform extraction followed by ethanol precipitation.

### 2.3. Reduced representation bisulfite sequencing (RRBS)

#### 2.3.1. Restriction enzyme digestion

A total of 3μg of high molecular weight genomic DNA was used for RRBS sample preparation. Each DNA sample was subjected to MspI restriction enzyme digestion. A total volume of 50μl was used in the procedure, including 3μl of MspI restriction enzyme (New England Biolabs) and 5μl of MspI reaction buffer (New England Biolabs). If the total volume was lower than 50μl, the difference was made up with nuclease-free water as recommended by the manufacturer. Incubation was performed in the thermocycler (Thermofisher) at 37°C for 15min. Next, the DNA purification procedure was performed using AmpureXp magnetic beads (Beckman Coulter).

#### 2.3.2. End repair

The DNA fragments with 5’-CG-3’ overhangs generated by the restriction enzyme digestion were end-repaired using Nextflex Bisulfite Kit (Bioscientific).

#### 2.3.3. Size selection, adenylation and adapter ligation

After end-repair, SPRI double size selection method combined with DNA purification was applied using AmpureXp magnetic beads (Beckman Coulter). Nextflex double size selection standard protocol was followed to select fragments between 200-300 bp (without adapters) with a mean length around 250 bp. A total volume of 20.5μl of preselected DNA was collected, and adenylation reactions were performed using adenylation mix, followed by incubation in the Thermocycler (Thermofisher) at 37°C for 30min. A different non-diluted adapter from Nextflex Bisulfite Barcodes Kit (Bioscientific) with a unique index sequence was chosen for each sample. Adapters were not diluted according to the manufacturer’s instructions. Ligation was performed for 15 min at 22°C. Subsequently, a DNA purification procedure was performed using AmpureXp magnetic beads (Beckman Coulter).

#### 2.3.4. Bisulfite conversion and amplification

The PCR products were purified using AmpureXp magnetic beads (Beckman Coulter) according to the Nextflex procedure. The purified fragments were then subjected to bisulfite conversion using the EZ DNA Methylation-Gold Kit (Zymo Research). The converted DNA was PCR amplified with some modifications. PCR reaction total volume was equal to 50μl, including 18μl of converted DNA, 22.75μl nuclease-free water, 2μl of Nextflex primer mix from Nextflex Bisulfite Barcodes Kit (Bioscientific), 1.25μl 10nM dNTP Mix (ThermoScientific), 5μl 1X Turbo Cx buffer (Agilent) and 2.5U Pfu Turbo Cx polimerase (Agilent). The thermocycling conditions: 2 min at 95°C and 12–18 cycles of 30 sec at 95°C, 30 sec at 65°C and 45 sec at 72°C, followed by a 7-min final extension at 72°C.

#### 2.3.5. Library validation and sequencing

The DNA libraries were quantified using the Qubit instrument (Life Technologies) and qualified using Agilent 2100 Bioanalyzer High Sensitivity chips (Agilent Technologies). According to the manufacturer’s instructions, paired-end sequencing (2× 100 bp) was performed on the Illumina HiSeq1500.

#### 2.3.6. Bioinformatics

Paired-end reads obtained from Illumina 1500 sequencer were filtered to remove the adapter contamination, and low-quality reads using the application TrimGalore!. Filtered reads were mapped to the reference genome (susScr3 version) using Bismark (Krueger and Andrews, 2011). Per CpG, methylation statistics were extracted using an application developed at the Department of Medical Genetics, Medical University of Warsaw. Positions with SNPs, changing CpG places to CpH or TpG, were also detected and filtered using the above application.

### 2.4. RRBS data analysis

Methylation levels of cytosines were analysed by methylKit.53 Briefly, the number of methylated and unmethylated CpG and non-CpG (CHG and CHH, H representing A/C/T) sites were counted for each region. CGIs were defined as regions >200 bp with a GC fraction >0.5 and an observed-to-expected ratio of CpG >0.6. CGI shores were defined as regions 2 kb in length adjacent to CGIs. To annotate porcine CGIs, reference genome (susScr3) and annotation were downloaded from USCSand and the Ensembl, respectively. To define the differentially methylated cytosines (DMCs), multiple pairwise comparisons were performed against CpG methylation information of twenty samples and filtered (q < 0.01) using methylKit.53

#### 2.4.1. Mapping

S.scrofa 10.2.79 and associated GTF files were downloaded from Ensembl. The fasta sequences were prepared for bismark (v0.14.3), them mapped using bowtie 1 as recommended for bismark software. To allow compatibility with bismark and methylkit, only somatic chromosomes were retained.

#### 2.4.2. Raw_reads and trimming

Raw reads were trimmed using TrimGalore (http://www.bioinformatics.babraham.ac.uk/projects/trim_galore/) with the following parameters -- trim1 --phred33 --length 50 --retain_unpaired –paired. Trimmed reads were mapped to the converted pig genome using bismark command bismark_v0.14.3/bismark --gzip -n 1 -1 pair1.fq.gz -2 pair2.fq.gz

#### 2.4.3. Identification of differentially methylated regions (DMRs)

MethylKit (https://code.google.com/p/methylkit/) was used to identify differentially methylated regions (DMRs) (bases and tiles), which show statistically significant differential methylation between two groups: high and low-efficiency AMSCs.

#### 2.4.4. annotation of genomic regions

High-density CpG promoter (HCP), intermediate-density CpG promoter (ICP), and low-density CpG promoter (LCP) annotations were defined based on the transcription start sites (TSS) of known RefSeq genes. In detail, HCP, which indicated the “CpG-rich” promoters, was identified as having a GC density ≥0.55 and the observed to expected CpG ratio (CpG O/E) ≥ 0.6; promoters with CpG O/E # 0.4 were classified as LCP; the remaining nonoverlapping promoter populations (0.4 < CpG O/E < 0.6) were classified as ICP. The annotated repeat elements such as LINEs, SINEs, and LTRs were downloaded directly from the RepeatMasker track of the UCSC Genome Browser. Other regions such as CGIs, exons, and introns were downloaded from the UCSC Genome Browser. Intragenic regions were included from the TSS to the transcription termination sites (TTS), whereas the intergenic regions were defined as the complement of the intragenic regions.

## 3. Results

### 3.1. Genomic location of Differentially methylated sites

We mapped the global DNA methylation patterns of adult male ADMSCs showing high and low efficiency of SCNT in pigs. We identified 3704 bases with statistically significant differences in methylation, 890 bases within 5Kb of a known transcript, 10062 tiles with statistically significant differences in methylation, and 4965 tiles within 5Kb of a known transcript.

### 3.2. Non-CG methylation: CHH, CHG

Fewer differentially methylated sites were seen in other contexts CHH 128 bases with statistically significant differences in methylation 39 bases within 5Kb of a known transcript, 2458 tiles with statistically significant differences in methylation 1354 tiles within 5Kb of a known transcript.

In the CHG context, we identified 59 bases with statistically significant differences in methylation, 23 bases within 5Kb of a known transcript, 2554 tiles with statistically significant differences in methylation and 1356 tiles within 5Kb of a known transcript.

### 3.3. Genome-wide CpG methylation and density patterns in relation to genomic features

Unsupervised hierarchical clustering of the individual methylation profiles of high and low efficient cells revealed separated sample groups (Figure 3). Thus, hierarchical clustering indicates that highly efficient and low efficient cells differ in methylation profiles.

**Figure 3.**
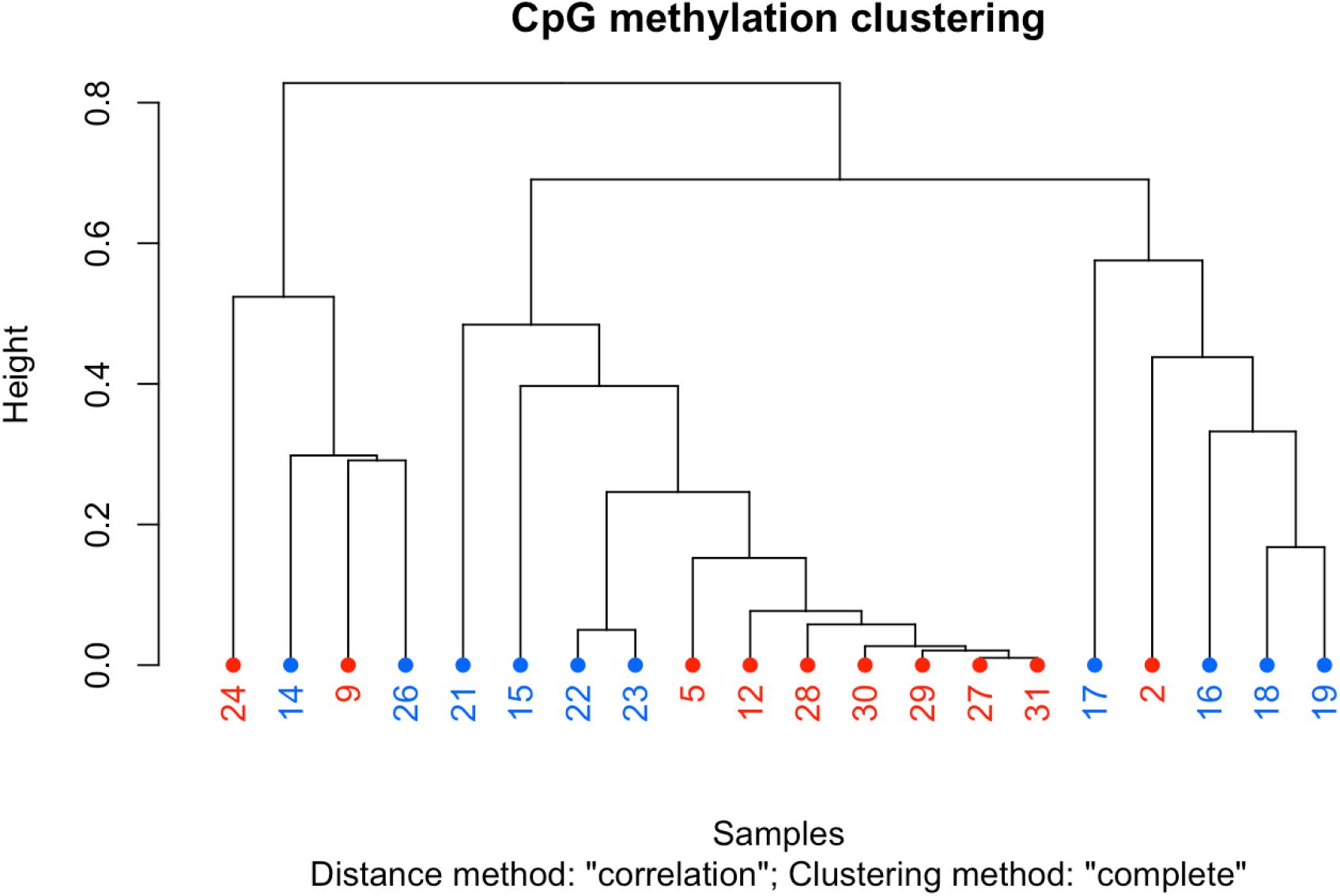
Hierarchical cluster analysis according to methylation patterns across analysed samples. Blue colour – high cloning efficiency cells; Red colour – low cloning efficiency cells. Dendrograms were produced using correlation distance and ward clustering methods. Numbers represent the individual sample ID.

## 4. Discussion

Cloning has become a powerful tool for analysing gene functions, genomic imprinting, genomic re-programming, development, neurodegenerative diseases, gene therapy, and more (Capecchi, 2000; Jang et al., 2010; Matoba and Zhang, 2018; Perrera and Martello, 2019). Somatic cell nuclear transfer is an essential cloning tool for biomedical and epigenetic research (Srirattana et al., 2022. This method enables significant development in biomedicine and disease modeling, where genome-edited mammals can be bred and used for disease research, transplantation, or to protect endangered species (Tian et al., 2003; Grzybek et al., 2017b; Fatira et al., 2018; Liu et al., 2021; Yue et al., 2021).

Due to its similarity to humans, the combination of somatic cell nuclear transfer (SCNT) and precise genome editing to generate transgenic pigs carrying required disease phenotype may be applied to swine (Wilmut et al., 2002; Yang et al., 2007). Here we showed that according to methylation patterns, there is a clear definition of groups with low and high cloning efficiency. Our study confirmed the presence of porcine CpG methylation patterns similar to those previously demonstrated for humans and mice and established the functional aspects of porcine CpG methylation (Ziller et al., 2011; Shirane et al., 2013; Schachtschneider et al., 2015).

DNA methylation in a promoter correlates with the transcription of a target gene (Niesen et al., 2005). Methylated genes are known to be linked with genomic region-specific DNA methylation patterns (Raza et al., 2017). We investigated promoter, exon and intron regions along the porcine genome and localised CpG islands to these genic features. The majority of differentially methylated sites were intergenic (Figure 2), and 37% were located in previously described CpG islands. We showed that methylation levels of CpG islands were lower than CpG island shores in the promoter, exon and intron regions. These results demonstrated that CpG islands located in different genic features displayed effects on the methylation patterns of the associated genes. A strong relation between methylations in CpG island shores located within 2 kb of an annotated transcription start site (TSS) and expression of associated genes was reported by Irizarry and others (Irizarry et al., 2009). CpG islands in exon regions showed different methylation levels than those in intron regions, suggesting that exons may affect the methylation patterns of CpG islands (Yuan et al., 2016; Chen et al., 2018).

Completion of the swine reference genome sequence (Groenen et al., 2012b; Groenen, 2016) gives a great ability to perform porcine studies for human diseases and disorders, as well as opens the door for targeted approaches to produce models for diseases (Gutierrez et al., 2015; Prather et al., 2015; Lunney et al., 2021). Our results provide novel information for future studies of the porcine epigenomics. The results based on RRBS are a powerful technology for epigenetic profiling of cell populations relevant to developmental biology and genetic engineering for porcine disease models. Further studies are necessary to investigate the similarities in methylation levels between humans and pigs for specific genomic regions. This knowledge will give a chance to analyse disease progression, the differences observed in intron and exon methylation patterns between pig tissues and human cell lines, and the proposed adaptive evolutionary role of CpG methylation.

In conclusion, porcine CpG methylation levels were similar to those reported for other mammals. We believe that our work will accelerate the practical use of the SCNT technique for pig model production and contribute to the studies of human disease, xenotransplantation, and molecular breeding in agriculture.

## Authors’ Contributions

The study was conceived and designed by MG & PL. Samples were maintained by PL. Laboratory work: PL, MG, MD, RP, KF. Data handling – MG. Statistical analysis was carried by TG, KF, PG and RDE. The manuscript was written by MG, MD, KF, TG. MG and MD revised the manuscript. Project administration – MG & PL. Funding acquisition – MG and PL.

